## Supplemental Figures for "Transposable element-mediated rearrangements are prevalent in human genomes"

SUPPLEMENTARY FIGURES

Supplementary Figure 1

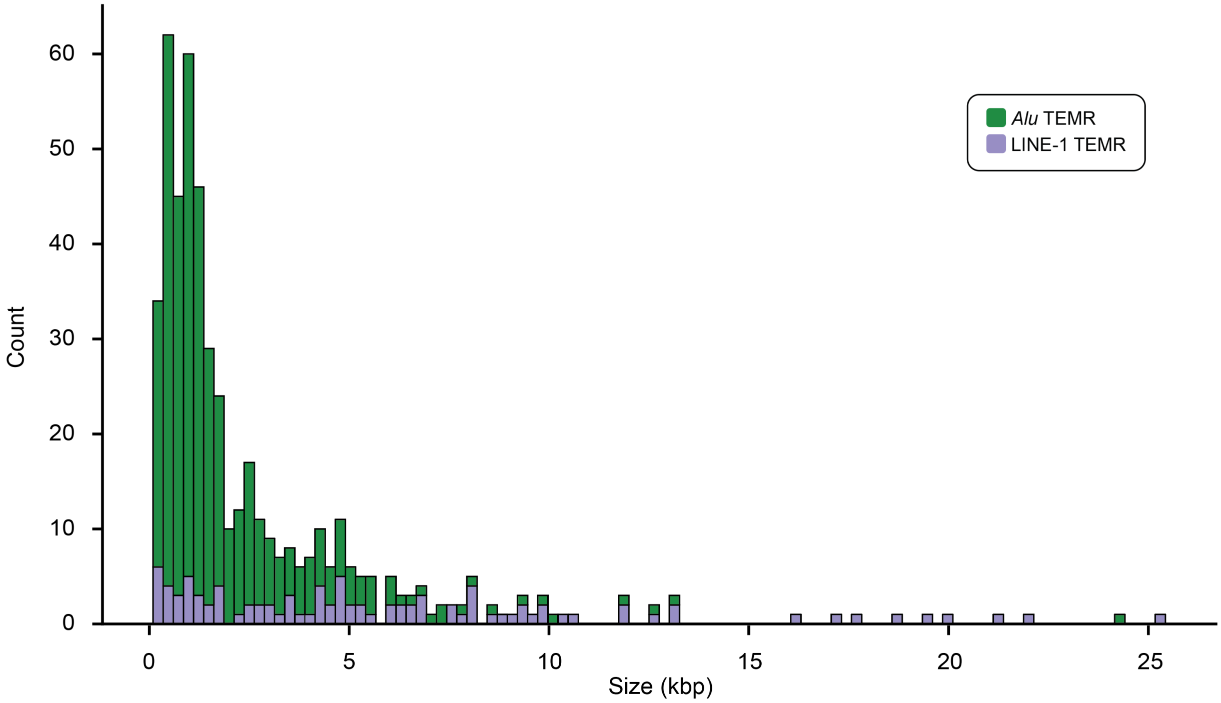


**Supplementary Figure 1.** Length distribution of *Alu* TEMR and LINE-1 TEMRs.

Supplementary Figure 2

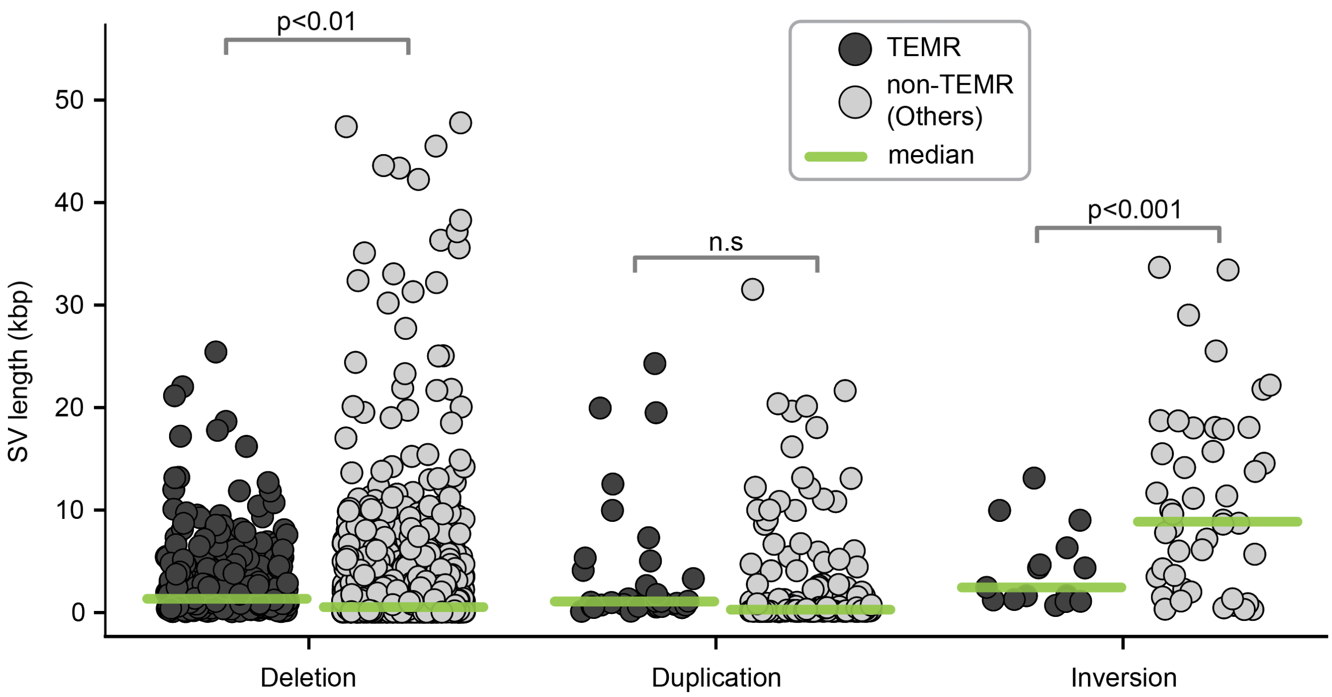


**Supplementary Figure 2.** Distribution of SV lengths between TEMR and non-TEMR rearrangements.

Supplementary Figure 3

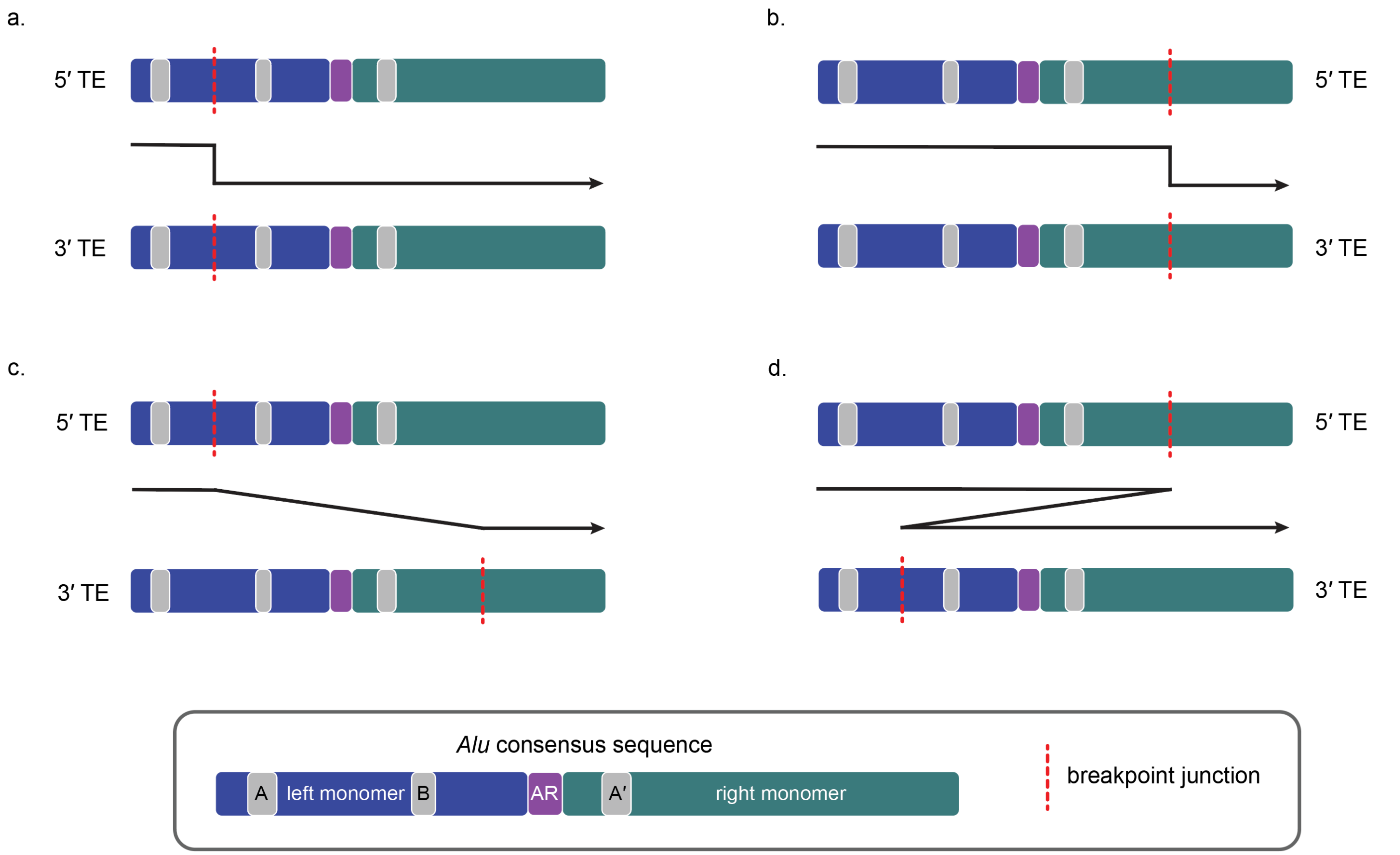


**Supplementary Figure 3.** Different recombination pattern between the two *Alu* elements from *Alu* TEMR-HR. Breakpoint junctions are present in the same monomer (a: left and b: right) of the two *Alu* elements resulting in a recombination product of ~300 bp chimeric *Alu*. Breakpoint junctions are present in the opposite monomer of the two *Alu* elements, resulting in a recombination product of either ~150 bp (c) or ~450 bp (d) chimeric *Alu*.

Supplementary Figure 4

a.

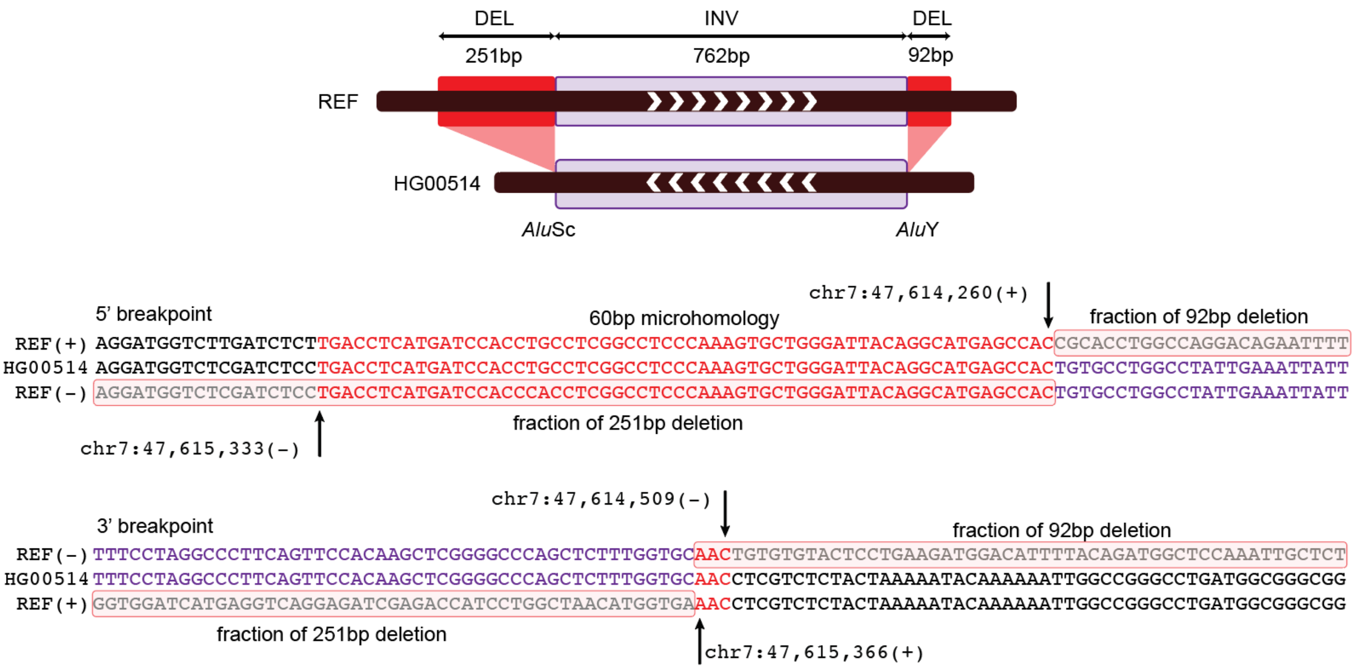


b.

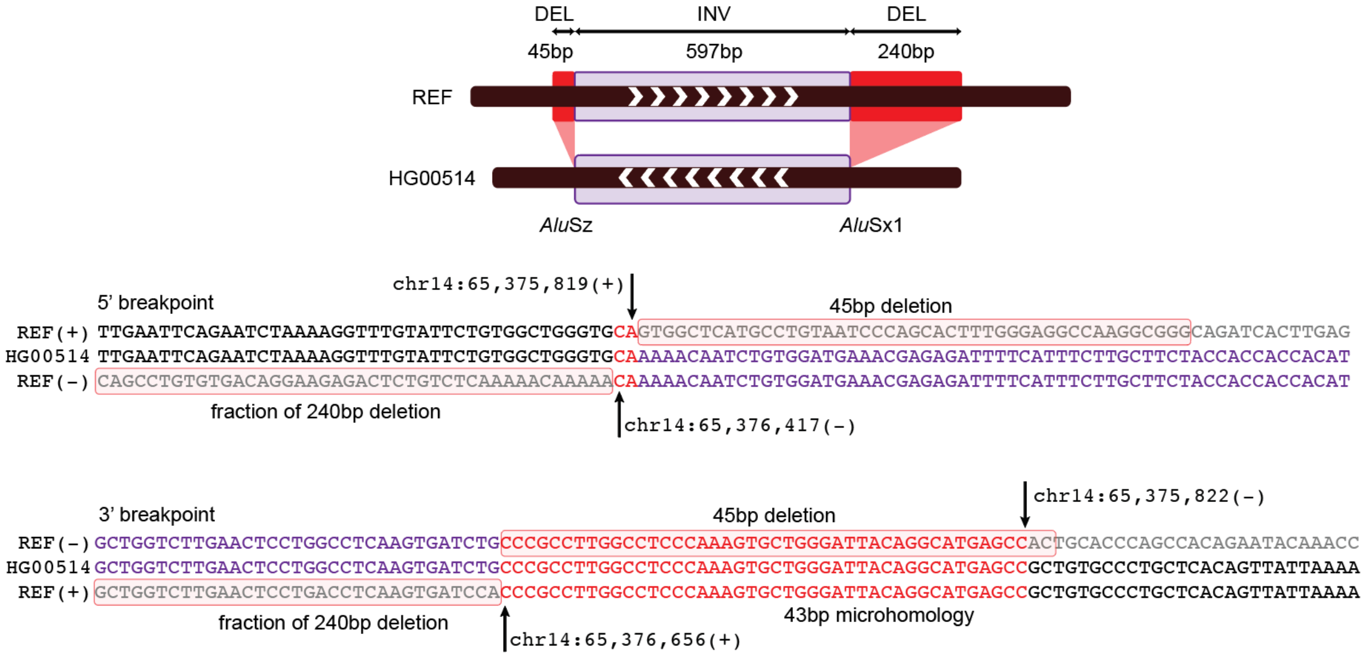


**Supplementary Figure 4. Inversions with complexities at the junctions**. a & b (top). Schematic representing the inversion (purple) with deletions (red) around the breakpoint junction. a & b (bottom) junction reconstruction of this complex inversion (purple font) event with their corresponding microhomologies (red font) and deleted sequence (red box).

Supplementary Figure 5

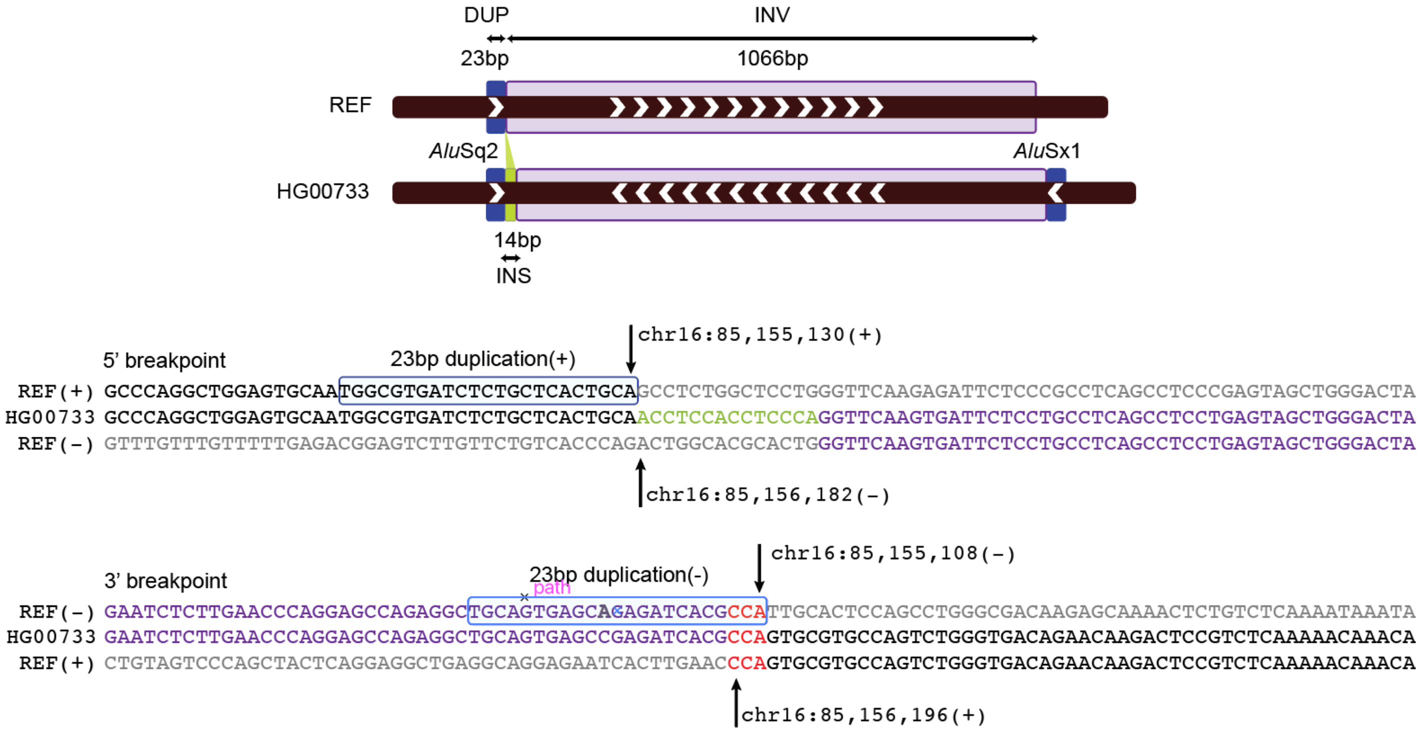


**Supplementary Figure 5. Inversions with complexities at the junctions.** Schematic (top) representing the inversion (purple) with accompanying duplication (blue) and insertion (green) deletions (red) around the breakpoint junction. Junction reconstruction (bottom) of this complex inversion (purple font) event with the corresponding microhomologies (red font), insertion (green font) and duplication sequence (blue box).

Supplementary Figure 6

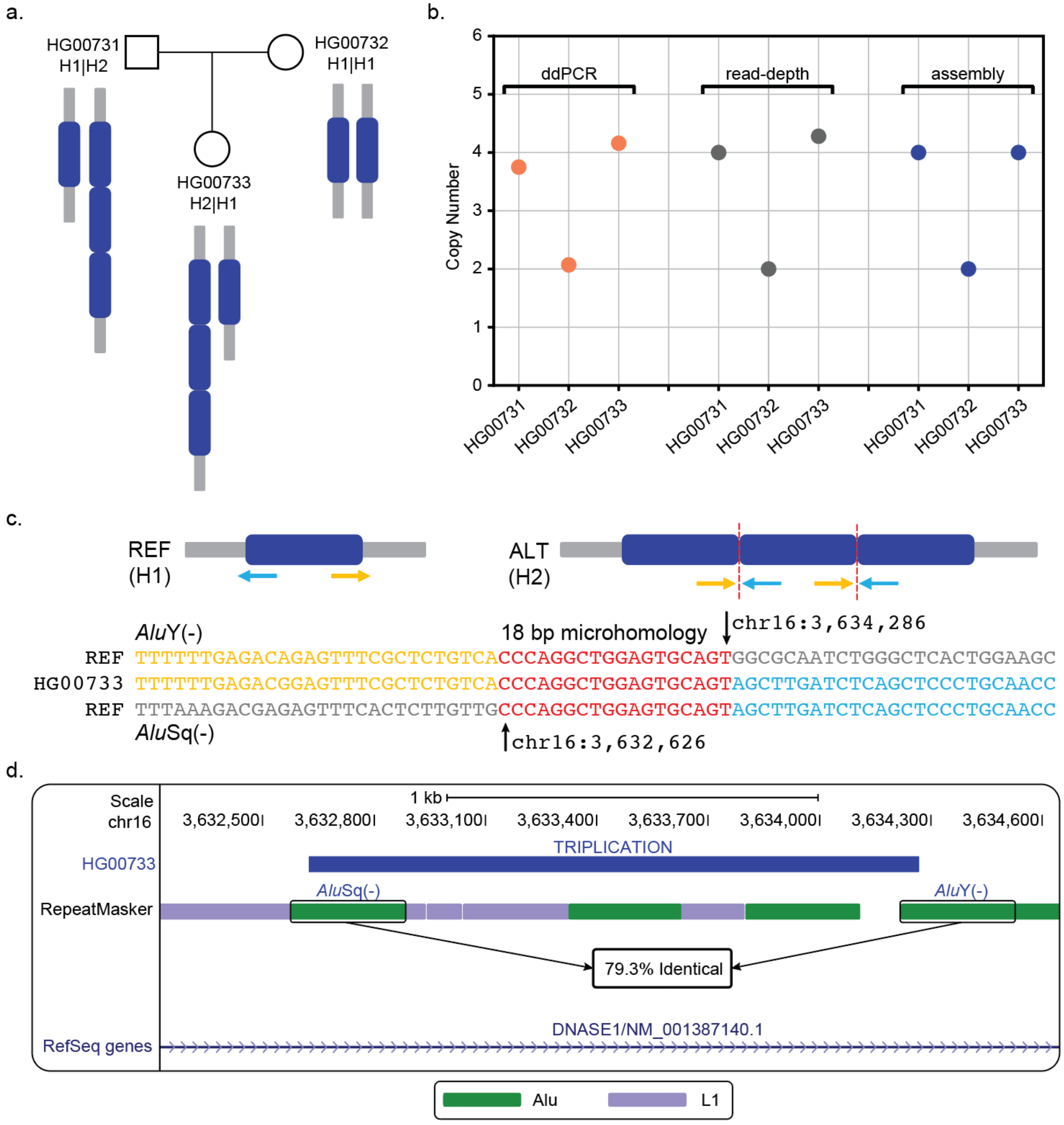


**Supplementary Figure 6. Complex rearrangements (mCNV) mediated by TEs. a,** Schematic representing the triplication found in the Puerto Rican trio. **b,** Copy number status of the individuals containing the triplication found using ddPCR, read-depth analysis and assembly data. **c,** Breakpoint junctions indicating an 18 bp microhomology. **d,** UCSC genome browser depicting the triplication between *Alu*Sq and *Alu*Y within the intron 1 of *DNASE1*.

Supplementary Figure 7

a.

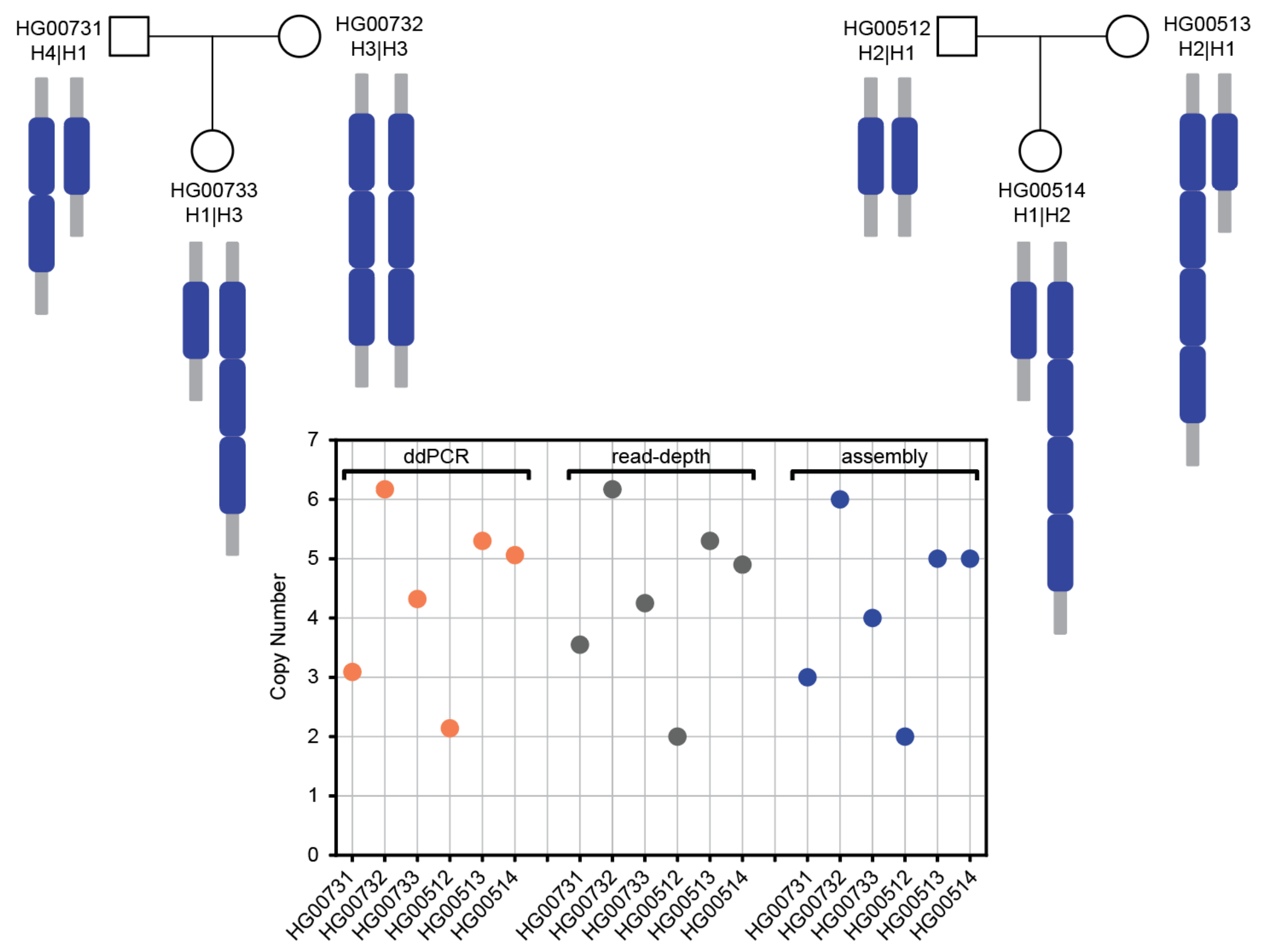


b.

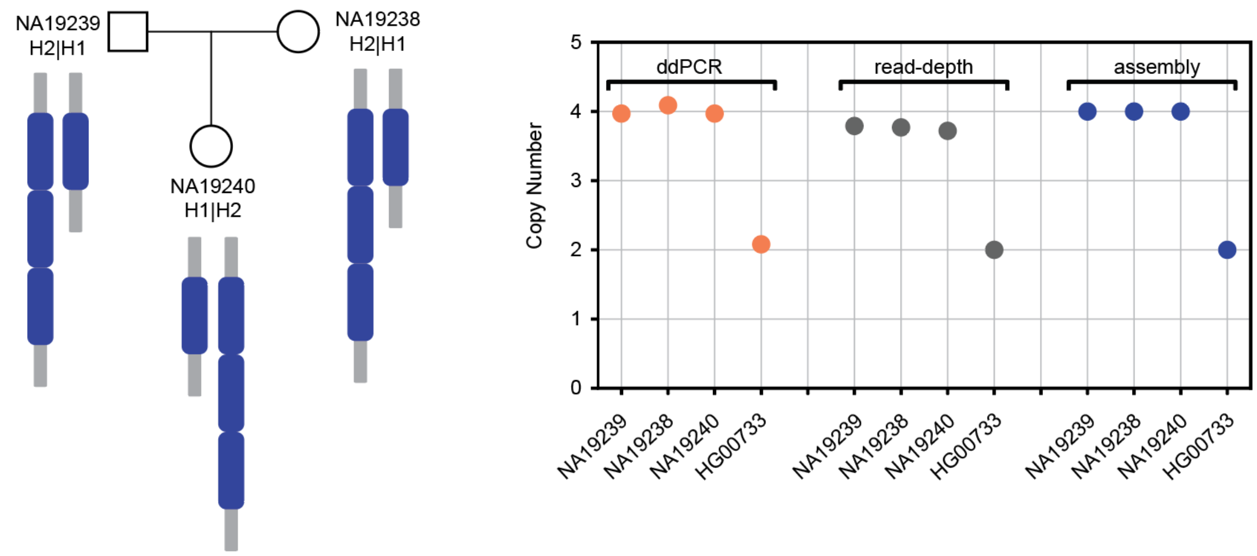


**Supplementary Figure 7. Complex rearrangements (mCNV) mediated by TEs. a,** Schematic representing the triplication (in Puerto Rican trio) and quadruplication (in Han Chinese trio) and copy number status of the individuals containing mCNV found using ddPCR, read-depth analysis and assembly data. **b,** Schematic representing the triplication found in the Yoruban (YRI) Nigerian trio and copy number status of the individuals containing triplication found using ddPCR, read-depth analysis and assembly data.

Supplementary Figure 8

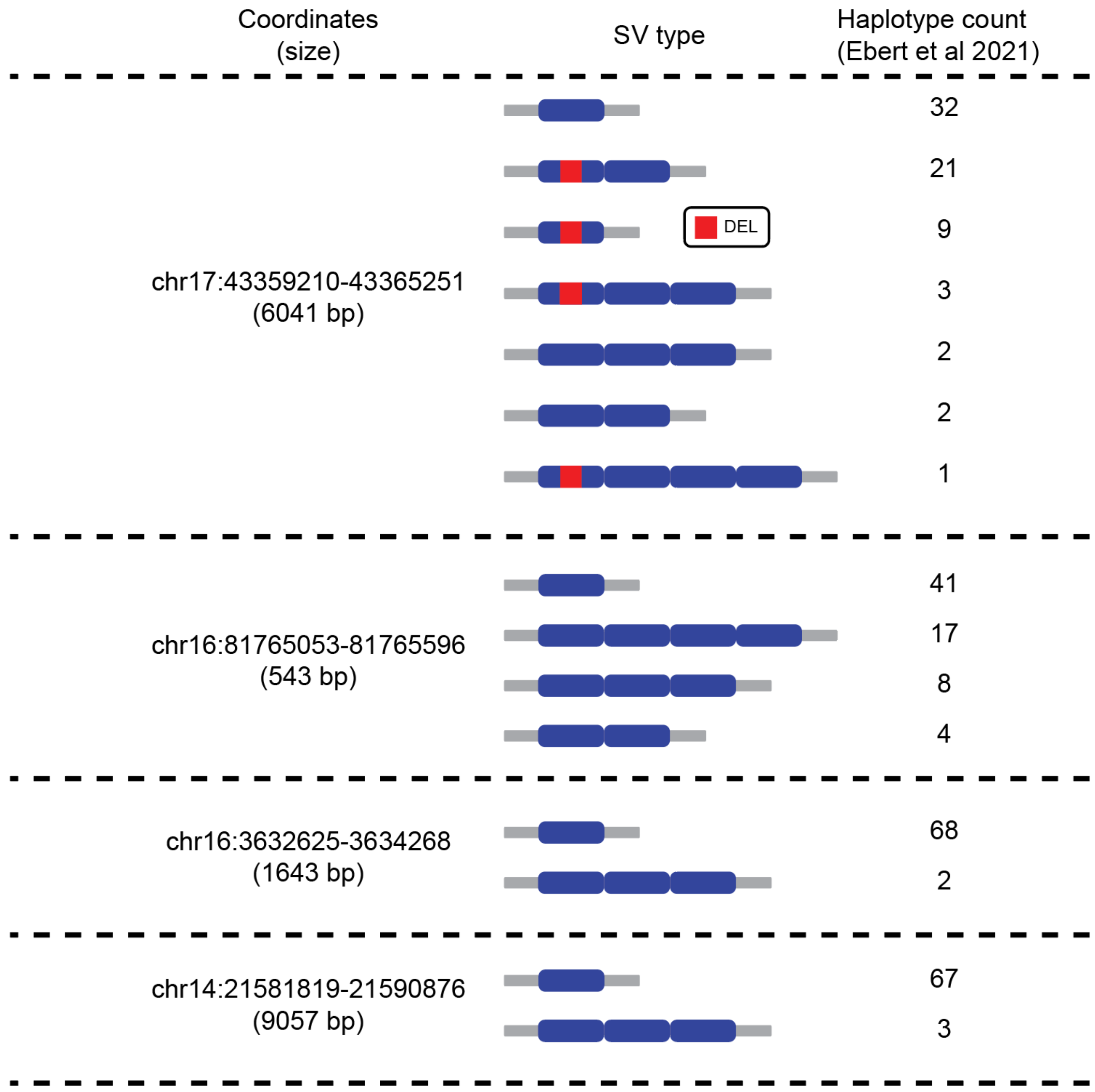


**Supplementary Figure 8.** Comparing four *Alu*/*Alu*-mediated amplifications with haplotype resolved human genomes from HGSVC.

Supplementary Figure 9

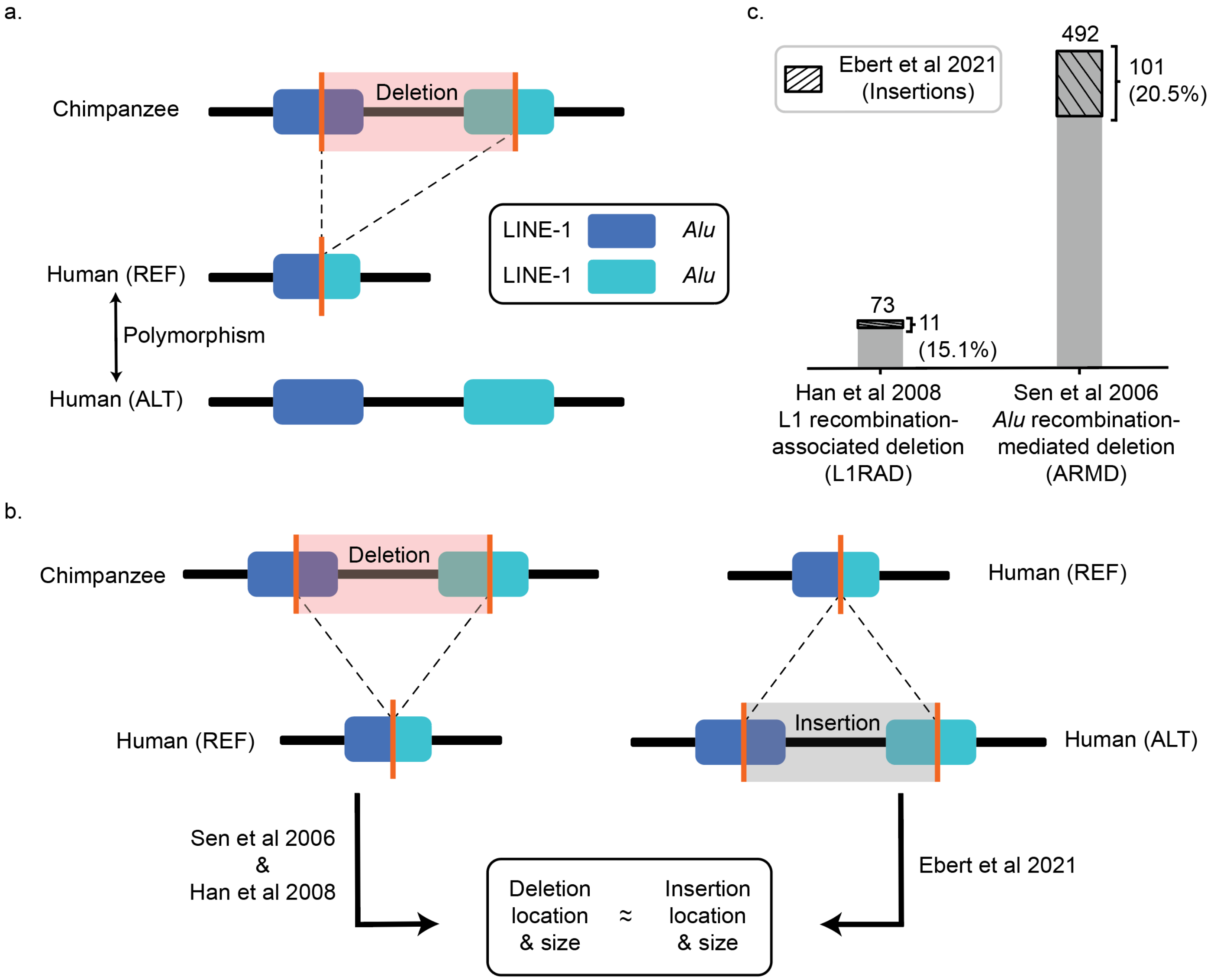


**Supplementary Figure 9. TEMR comparison between human and chimpanzee. a,** Schematic representing polymorphic TEMRs between human and chimpanzee genomes. **b,** Schematic representing the method used to compare outputs between studies. **c,** Comparing results between studies and identifying the percentage of TEMRs that are polymorphic within humans.

Supplementary Figure 10

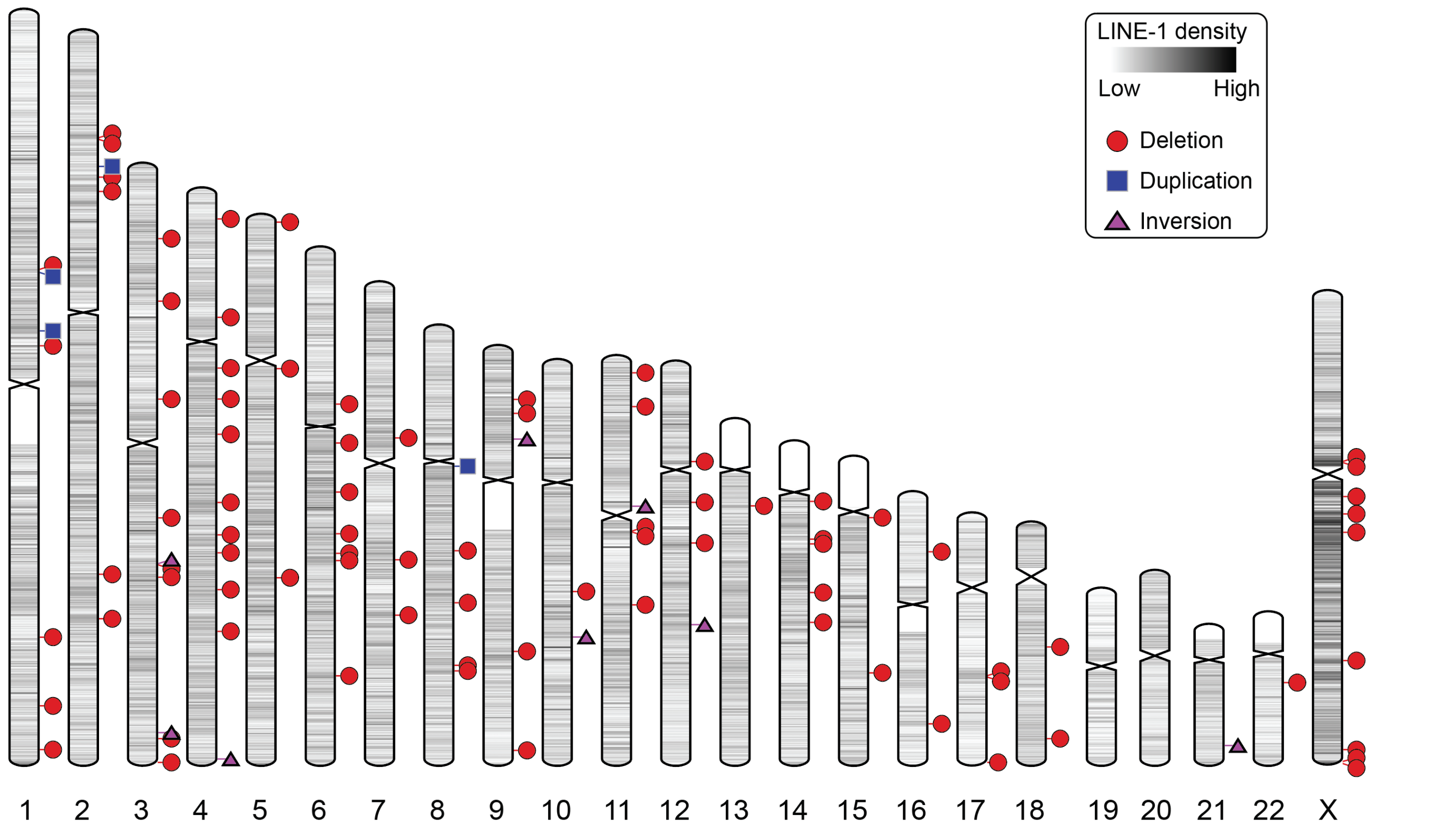


**Supplementary Figure 10.** Ideogram displaying 96 LINE-1 TEMRs. The intensity of shade represents the level of LINE-1 density.

Supplementary Figure 11

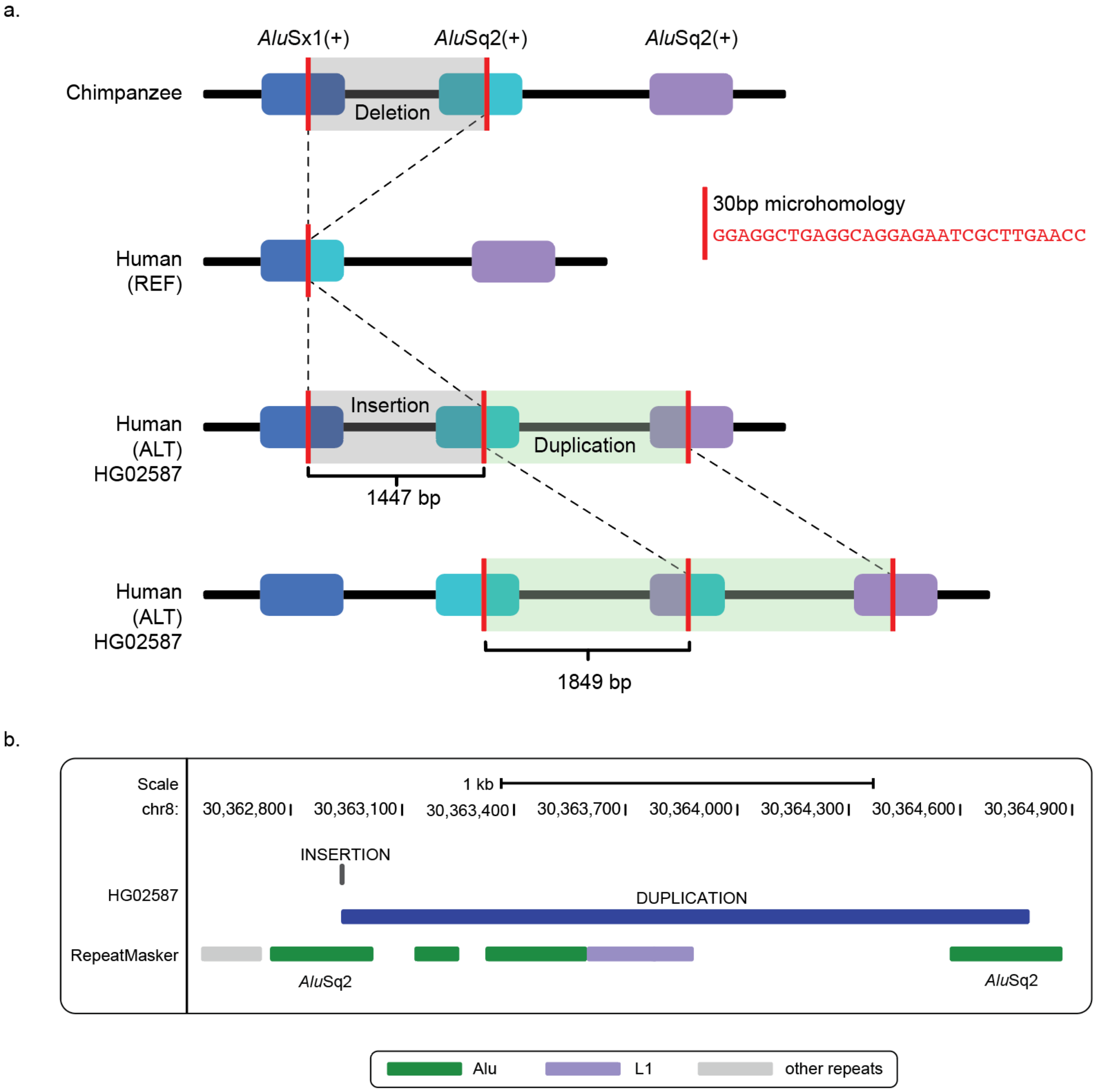


**Supplementary figure 11. Example of a single *Alu* being involved in two separate rearrangements. a,** Polymorphic deletion between *Alu*Sx1(blue) and *Alu*Sq2(cyan) and a duplication between *Alu*Sq2 (cyan) and *Alu*Sq2 (purple) in HG02587. **b,** UCSC genome browser depicting the insertion within *Alu*Sq2 and duplication between *Alu*Sq2 and *Alu*S2.

Supplementary Figure 12

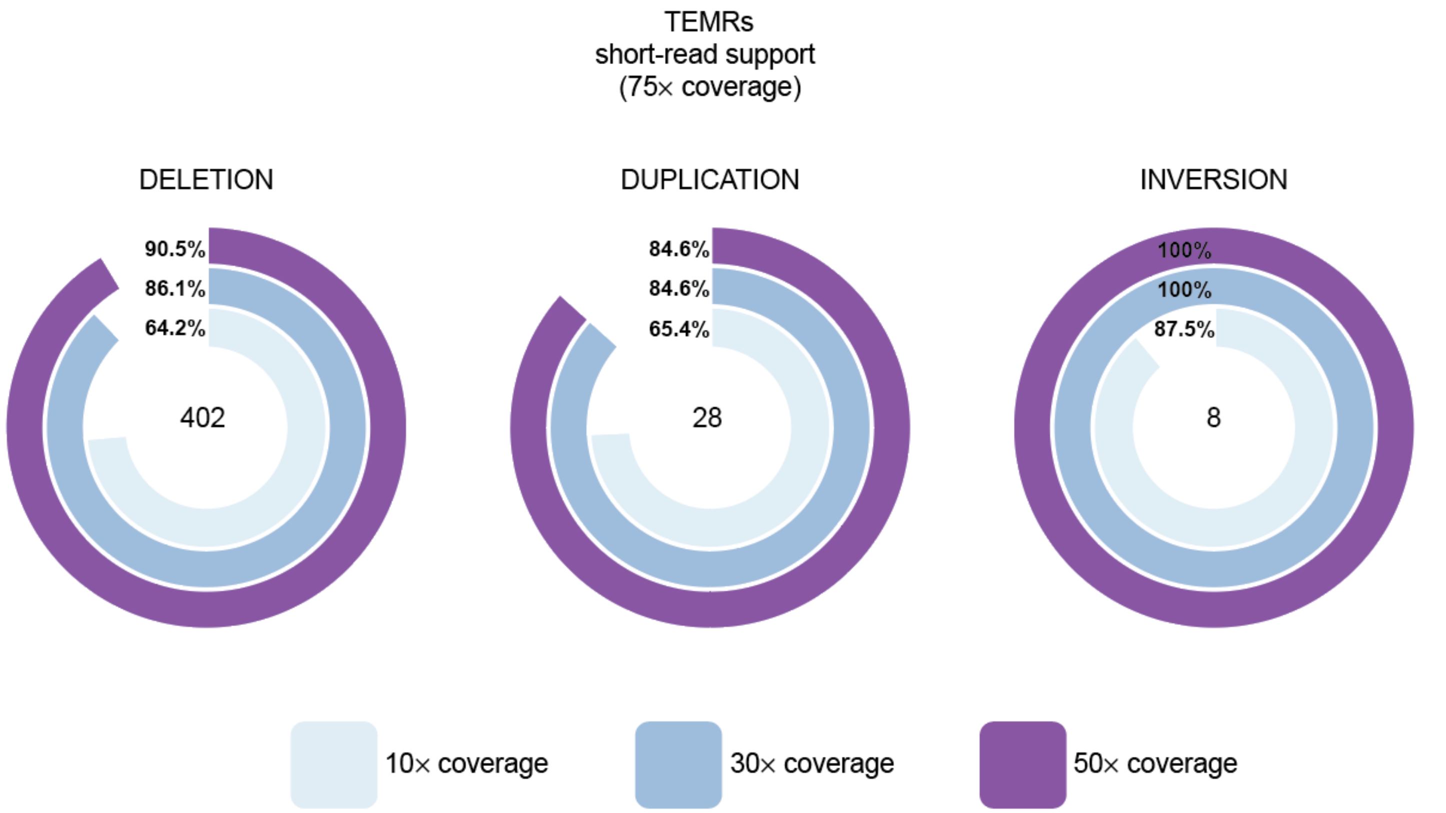


**Supplementary Figure 12.** Percentage of TEMRs identified using lower coverage (down sampled to 50×, 30× and 10×) short-read HTS data compared to original (75×) short-read HTS data used in this study.

Supplementary Figure 13

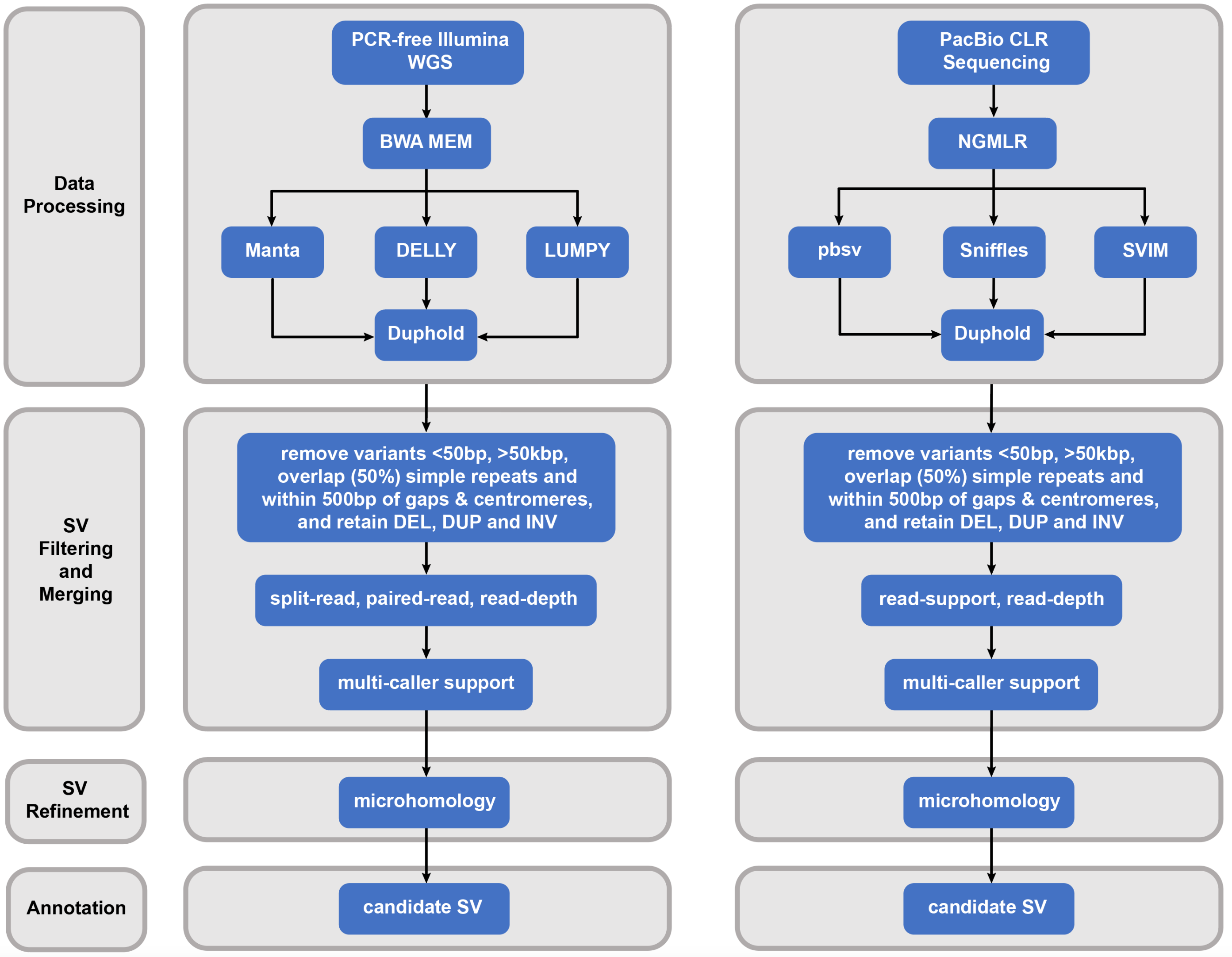


**Supplementary Figure 13.** Pipeline for identifying non-redundant high-confident SVs using short-read and long-read HTS data.

Supplementary Figure 14

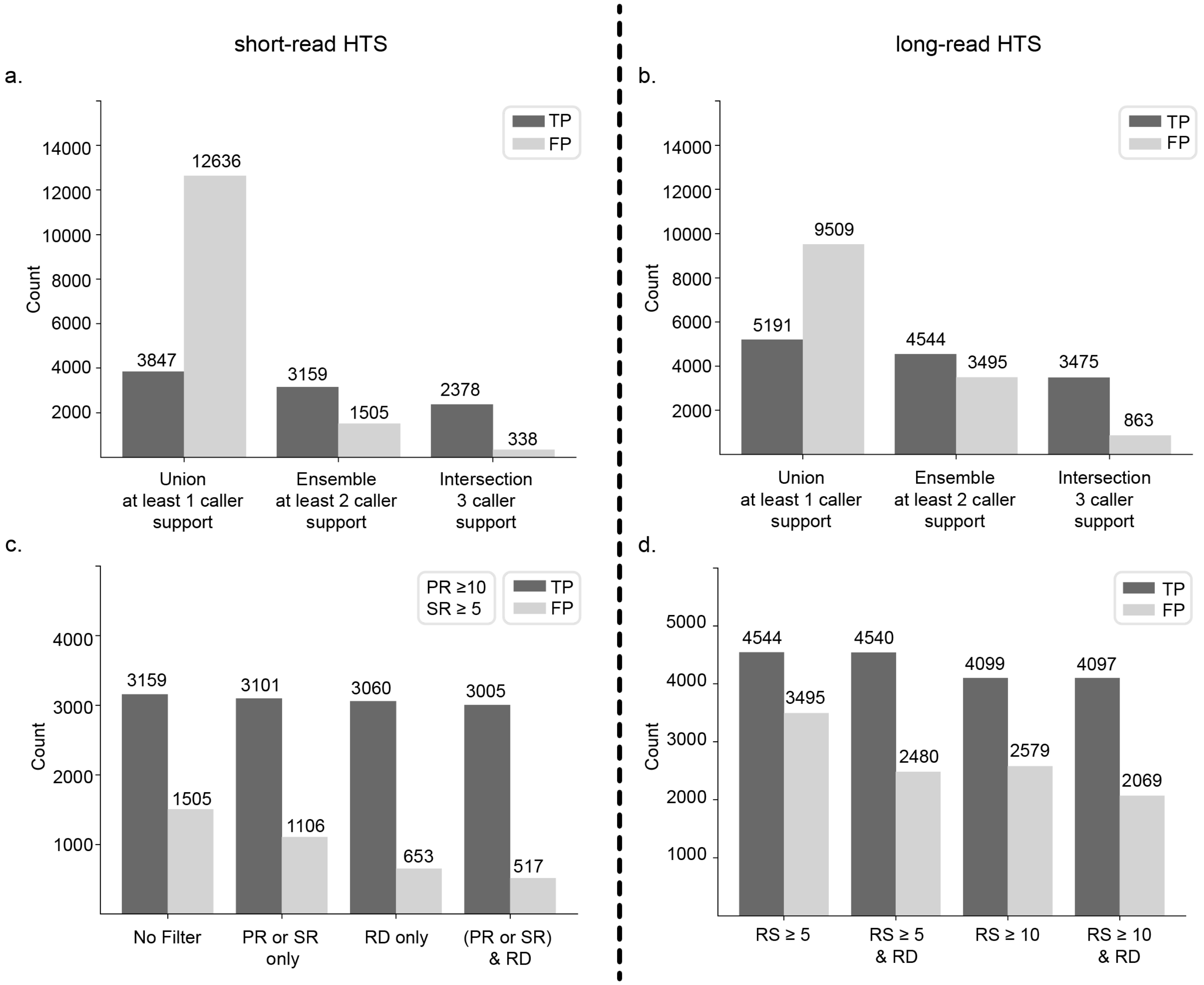


Supplementary Figure 14. Different approaches used for merging and filtering SVs using short-read HTS (a, c) and long-read HTS (b, d) data. **a, b**. Merging data from multiple callers. Average true positive SVs (TP: dark gray) compared to the average false positive SVs (FP: light gray) per sample, for methods with at least 1 (Union) or 2 (Ensemble) or 3 (Intersect) caller support for identifying an SV. **c, d**. Different parameters (paired-read: PR, split-read: SR, read-depth: RD) used for additional filtering. Average true positive SVs (TP: dark gray) compared to the average false positive SVs (FP: light gray) per sample.

Supplementary Figure 15

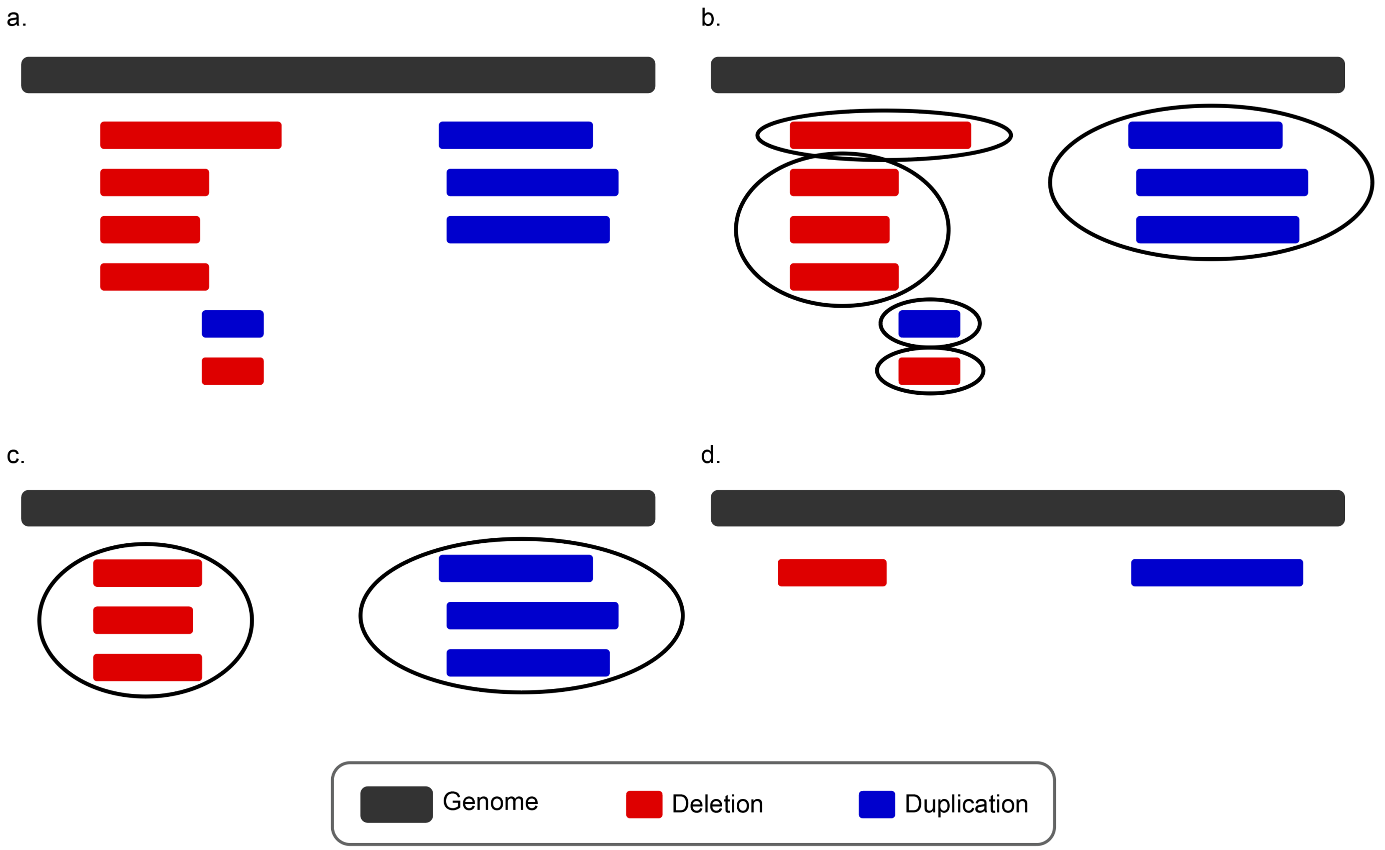


Supplementary Figure 15. Rank-based merging SVs. **a,** SVs are sorted based on coordinates. **b,** SVs are clustered by 80% reciprocal overlap and type. **c,** Clusters without SVs from at least 2 callers are removed. **d,** SV with the highest rank within each cluster is retained and the rest are removed, resulting in 1 SV per cluster.
